# Single-cell digital twins identify drug targets and repurposable medicine in Alzheimer’s disease

**DOI:** 10.1101/2025.01.14.633081

**Authors:** Yunxiao Ren, Ming Hu, Yang E. Li, Andrew A. Pieper, Jeffrey Cummings, Feixiong Cheng

## Abstract

Alzheimer’s disease (AD) is a complex and poorly understood neurodegenerative disorder without sufficiently effective treatments. Novel approaches to identify FDA-approved drugs that may hold potential for mitigating symptoms of AD hold promise for addressing this problem. One such strategy is the use of digital twins (DTs), which are virtual representations of physical entities that facilitate therapeutic target identification by accurately characterizing disease heterogeneities in real time through continuous feedback and dynamic model updates. In this study, we developed a single-cell digital twin (scDT) framework using single-nuclei RNA-seq (1,197,032 nuclei) and ATAC-seq (740,875 nuclei) data from the middle temporal gyrus of 84 donors across 4 degrees of AD neuropathological change (ADNC). We observed differential gene expression for six major cell types intensified at severe ADNC. We also constructed cell type-specific transcription factor (TF)-target gene networks by leveraging peak-to-gene linkages and motif enrichment analyses. By integrating genome-wide association study (GWAS) loci with cell type-specific *cis*-regulatory DNA elements (CREs), we identified 141 ADNC-associated genes. Using gene set enrichment analysis (GSEA) and network proximity analysis, we identified 13 candidate repurposable drugs that were associated with these ADNC-associated genes. In summary, we constructed a single-cell digital twin (scDT) framework cell type-specific target identification and drug repurposing in AD and other complex diseases if broadly applied.

## 1. Introduction

Alzheimer’s disease (AD) is a complex, age-related neurodegenerative disorder and the leading cause of dementia worldwide^1^. The accumulation of amyloid-β (Aβ) plaques and tau protein tangles is a hallmark of AD pathology. However, it is increasingly recognized that additional neuropathological processes, such as neuroinflammation, lipid metabolism dysfunction, mitochondrial impairment, axonal degeneration, impaired postnatal neurogenesis, and synaptic dysfunction, also play crucial roles in disease progression^1–3^. This intricate pathology poses significant challenges for developing effective therapies. Recent advances in single-cell multi-omics have illuminated cell type-specific dynamic changes of transcriptome and epigenome in AD heterogeneities and disease progression^2,4–10^, revealing new complexities in disease mechanisms. Furthermore, intercellular heterogeneity complicates therapeutic efforts, and substantial differences in the molecular alterations between laboratory animal models and human patients highlighting the limitations of traditional laboratory-based approaches to therapeutic discovery^11–13^. Given the magnitude of the problem, innovative strategies for discovering treatments for AD are essential.

The concept of the digital twin (DT), inspired by the industrial sector, represents a virtual model of a cellular system or organ system, or even the entire human body^14–19^. DTs have transformative potential for biomarker identification, drug discovery, and drug repurposing by facilitating bidirectional interactions between physical and digital realms, paving the way for precision medicine^20–23^. By integrating multi-dimensional datasets to create highly specific virtual representations, DTs leverage advanced technologies such as generative artificial intelligence (AI), statistical modeling, network medicine, and multi-omics integration^24,25^. With the escalating burden of AD, research efforts have generated extensive datasets through large-scale initiatives^10,26–30^. By harnessing these datasets, DTs have potential to accelerate therapeutic advancements and enhance our understanding of AD pathophysiology.

In this study, we developed a single-cell Digital Twin (scDT) framework to model AD progression, enabling therapeutic target identification and potential drug repurposing. Using single-nuclei RNA-seq (snRNA-seq) and ATAC-seq (snATAC-seq) data from the middle temporal gyrus of 84 donors across four stages of AD neuropathological changes (ADNC): non-AD, low, intermediate (Inter), and high ADNC, we provide stage-specific transcriptomic and epigenomic profiles for six major brain cell types throughout ADNC. Through constructing cell type-specific transcription factor (TF)-target gene regulatory networks, we also identified cell type-specific candidate *cis*-regulatory DNA elements (cCREs), shedding light into the regulatory mechanisms underlying ADNC. By integrating genome-wide association study (GWAS) loci with cCREs, we then identified 141 AD-associated genes. This provided a basis for exploring drug repurposing through gene set enrichment analysis (GSEA) and network proximity approaches, which identified 13 candidates for AD treatment (galantamine, mecamylamine, dextromethorphan, tubocurarine, gabapentin, ebselen, resveratrol, imatinib, dipyridamole, deferoxamine, gemfibrozil, lansoprazole, and probucol). Thus, our findings demonstrate the utility of single-cell DTs in unraveling the complexity of AD progression, providing a potential platform for cell type-specific therapeutic targets identification and drug repurposing for AD and other neurodegenerative diseases if broadly applied.

## 2. Materials and Methods

### 2.1. Resources of snRNA-seq and snATAC-seq data

The snRNA-seq and snATAC-seq data used in this study are available from the Seattle Alzheimer’s Disease Brain Cell Atlas (SEA-AD) consortium (syn26223298)^10^. We obtained the gene expression count matrices in the h5ad format and the snATAC-seq raw fragments files from 84 human brain middle temporal gyrus samples. We also acquired the corresponding metadata manifest (Supporting Information **Table S1**), which includes comprehensive neuropathological staging metrics such as Thal phase, Braak stage, Consortium of Establish a Registry of Alzheimer’s Disease (CERAD) score, and overall AD neuropathological change (ADNC). The metadata also provides demographic and clinical information, including sex and the presence of co-pathologies such as Lewy Body Disease and Limbic-predominant age-related TDP-43 encephalopathy (LATE).

### 2.2. snRNA-seq data quality control, dimensionality reduction and clustering

The snRNA-seq count matrices were processed using the Scanpy (v1.9.5) package^31^ in a python 3.10 environment. Quality control metrics were initially computed using the calculate_qc_metrics function. Genes expressed in fewer than 10 cells and cells with fewer than 200 detected genes were excluded of the analysis. Outliers for total_counts and n_genes_by_counts were identified using the interquartile range (IQR) method. First, the 75th percentile (Q3) and 25th percentile (Q1) of the respective fields were calculated. The IQR was then determined as: *IQR = Q3 − Q1*. Based on this, lower and upper thresholds were defined as: *Lower threshold = Q1 – 3 × IQR*; *Upper threshold = Q3 + 3 × IQR*. Values outside this range were flagged as potential outliers, and only cells with values falling within the lower and upper thresholds were retained. For mitochondrial (pct_counts_mt) and ribosomal (pct_counts_rb) percentages, outlier thresholds were determined by combining the 3×IQR rule with fixed cutoffs. The upper threshold was computed using the 3×IQR rule: Upper threshold (IQR) = Q3 + 3 × IQR. This threshold was then compared with fixed cutoffs of 10% for pct_counts_mt and 20% for pct_counts_rb. The final upper threshold was the maximum of the IQR-based threshold and the fixed cutoff. Cells with values exceeding this final upper threshold were classified as outliers and excluded from further analysis. Finally, doublets were identified and excluded using the Scrublet tool^32^, further refining the dataset for downstream analyses.

We normalized the processed count matrices using a shifted logarithmic transformation to stabilize variance^33^. Principal components (PCs) were computed to reduce dimensionality, and the data were clustered using the Leiden algorithm^33,34^. To visualize the clustering results, we utilized the Uniform Manifold Approximation and Projection (UMAP) technique^35^. Major cell type annotations for each cluster were assigned based on established marker genes^5,8,10,36,37^ (Supporting Information Fig. S1). We also compared these annotations with those reported in the original paper and found them to be consistent^10^. Neuron cell subtypes were labeled according to the classifications in the original paper_10_.

### 2.3. Differential expression gene (DEG) and gene functional enrichment analysis

We employed the R package muscat to identify differentially expressed genes (DEGs) by aggregating single-cell data into pseudobulk profiles^38^. Each gene was tested for expression changes within individual clusters, resulting in a total of “#genes × #clusters” tests for each comparison of interest^38^. Specifically, we focused on condition-related changes across AD progression stages, analyzing six comparison groups: Low-AD versus (vs.) Non-AD, Inter-AD vs. Non-AD, High-AD vs. Non-AD, Inter-AD vs. Low-AD, High-AD vs. Low-AD, High-AD vs. Inter-AD. DEGs were defined as genes meeting the criteria of a *p*-value < 0.05 and an absolute log fold change (|logFC|) > 0.5, ensuring statistical significance and biologically meaningful changes.

Gene functional enrichment analysis was conducted using the clusterProfiler package (v4.10.1)^39–41^. Gene Ontology (GO) enrichment was evaluated for terms related to Biological Process (BP) and Molecular Function (MF) across different stages of AD progression for each cell type. Enrichment results were filtered using the Benjamini-Hochberg (BH) method (adjusted *p*-value < 0.05).

### 2.4. snATAC-seq data quality control, dimensionality reduction and clustering

The snATAC-seq fragment data were processed using the ArchR R package (v1.0.3)^42^ in an R 4.4.2 environment, employing the hg38 reference genome for annotation. For initial quality control, three key metrics were assessed: the number of unique nuclear fragments, the signal-to-background ratio calculated by transcription start site (TSS) enrichment score, and the fragment size distribution^42^. To ensure sufficient data for reliable analysis, cells with fewer than 1,000 fragments were excluded (minFrags = 1000). The signal-to-background ratio was quantified using TSS enrichment score, with a minimum threshold of 4 (minTSS = 4). Cells not meeting these criteria were filtered out. To further enhance data quality, doublets were identified and removed using the ArchR functions addDoubletScores and filterDoublets, with a filter ratio of 1.

Dimensionality reduction was performed using the iterative Latent Semantic Indexing (LSI) approach implemented in the ArchR package via the ‘addIterativeLSI’ function^42^. The process utilized three iterations (iterations = 3), with the final iteration sampling 30,000 cells (sampleCellsFinal = 30, 000). All other parameters are set to their default values. Clustering was performed using the ‘addClusters’ function in ArchR, initially with a resolution of 1 and finalized with a resolution of 0.4. Clusters containing fewer than 100 cells were excluded as low-quality clusters. Marker genes for each cluster were identified using the ‘getMarkerFeatures’ function, which uses the Gene Activity Score (accessibility around each gene) as a proxy for gene expression^43^. Cell types were annotated based on established marker genes (Supporting Information Fig. S1). Gene activity scores were visualized by UMAP, with smoothing applied using the MAGIC algorithm^44^ to enhance visualization.

### 2.5. Peak calling and annotation

Peak calling was performed using MACS2^45^ with default parameters. Cell type-specific marker peaks were identified using the ‘getMarkerFeatures’ function in the ArchR package. CREs specific to each cell type were further characterized by integrating these marker peaks with annotations from the hg38 reference genome. Transcription factor (TF) motif enrichment within these marker peaks was analyzed using the cisBP database^46^ through the ‘addMotifAnnotations’ and ‘peakAnnoEnrichment’ functions, specifying peakAnnotation = ‘Motif’. Additionally, footprinting analysis for motifs of interest was conducted using the ‘getFootprints’ function in ArchR^42^, providing insights into motif accessibility and regulatory activity.

### 2.6. Integration of snRNA- and snATAC -seq datasets

We integrated snATAC-seq data with corresponding snRNA-seq data for each stage of AD progression, following previously established methods^43^. This integration was performed using the ‘addGeneIntegrationMatrix’ function in the ArchR package, which leverages Seurat’s ‘FindTransferAnchors’ function to align datasets through canonical correlation analysis (CCA). For the integration process, we specified nGenes = 2,000 and dimsToUse = 1:40. Initially, we conducted unconstrained integration by setting addToArrow = FALSE to create a preliminary alignment. We subsequently refined the integration results by applying a constrained approach.

### 2.7. Linked AD associated GWAS loci with cCRE

We obtained GWAS summary statistics from the study by Bellenguez et al^47^ to identify AD-associated loci. Significant AD-associated GWAS loci were identified by applying a *p*-value threshold of *p*-value < 5e8. Additionally, We got AD lead SNPs of this study^47^ from https://www.ebi.ac.uk/gwas/studies/GCST90027158, which excludes 23andMe samples. In total, we identified 88 lead SNPs. We also identified cell type-specific CREs from the significant cell marker peaks (FDR ≤ 0.05 and Log2FC > 0.5). To establish functional links between these AD-associated loci and cCREs, we determined overlaps between the genomic positions of the AD-associated loci and CREs.

### 2.8. Identification of positive TF-regulators and putative targets

TFs can be identified based on changes in chromatin accessibility at DNA-binding motif sites using ATAC-seq. However, precise identification is often hindered due to the similarities of binding motifs among certain TF families by position weight matrices (PWMs)^42^. To address this, we integrated gene expression data to identify TFs with expression that correlated positively with chromatin accessibility changes using ArchR package as described previously^42,43^. This method correlates chromVAR deviation z-scores of TF motifs with the gene expression of corresponding TF across low-overlapping cell aggregates. Initially, deviant TF motifs were identified by calculating the maximum delta in deviation z-scores across clusters, enabling the stratification of motifs based on inter-cluster variability. We then employed the ‘correlateMatrices’ function in ArchR to compute the correlation between the gene expression matrix and the motif matrix. These correlations are assessed across many low-overlapping cell aggregates identified in the lower dimension space^42^. Positive TF regulators were defined based on the following stringent criteria: a correlation coefficient (cor) between motif activity and gene expression greater than 0.5, an adjusted *p*-value below 0.01, and a maximum inter-cluster z-score variation exceeding 0.75.

We identified the regulatory targets of these positive TF-regulators following the methodology described in the previous study^43^. First, we calculated the Pearson correlation coefficient between the chromVAR motif activity of positive TF regulators and the integrated expression levels of all expressed genes. We then computed the linkage score (LS) for each gene-TF pair using the following equation:

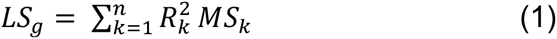

Here, LSg: Linkage score for gene g. n: Number of peaks linked to gene g. Rk: Pearson correlation coefficient between peak k and gene g. MSk: Motif score for the TF motif present in peak k. This scoring method ensured that higher linkage scores indicate stronger regulatory relationships, reflecting a greater number of linked peaks containing the TF motif, as well as strongly correlated peaks enriched for the motif.

Putative target genes for positive TF-regulators were filtered based on an LS threshold set at the 80th percentile of all LS values, retaining genes with LS values exceeding this threshold. Additionally, only genes with a correlation coefficient > 0.5 and FDR < 0.05 were considered. GO term enrichment analysis of the putative target genes was subsequently performed using the topGO package^48^.

### 2.9. Building human protein–protein interactome and drug–target network

The human protein-protein interactome (PPI) and drug-target network were constructed in our previous studies^46–50^. In this study, the human PPI was constructed by integrating data from multiple established PPI databases supported by experimental evidence. This includes binary PPIs identified by high-throughput yeast-two-hybrid (Y2H) experiments^54^ and kinase-substrate interactions derived from both low- and high-throughput literature sources, including the Human Protein Resource Database (HPRD)^55^, dbPTM 3.0^56^, Phospho.ELM^57^, KinomeNetworkX^58^, PhosphoNetworks^59^, and PhosphositePlus^60^. Additionally, signaling networks were curated from literature-based low-throughput experiments provided by SignaLink2.0^61^. Protein complex interactions, comprising approximately 56,000 candidate interactions, were identified using robust affinity purification-mass spectrometry data from BioPlexV2.0^62^. Furthermore, literature-curated PPIs derived from affinity purification coupled with mass spectrometry were included from sources such as HPRD^63^, PINA^64^, MINT^65^, InnateDB^66^, IntAct^67^, Instruct^68^, and BioGRID^69^. In total, this dataset encompasses 351,444 PPIs involving 17,706 protein nodes. The complete dataset is publicly available at https://alzgps.lerner.ccf.org.

The drug–target network was developed by integrating multiple reputable data sources, including DrugBank (version 4.3)^70^, BindingDB^71^, ChEMBL (version 20)^72^, the Therapeutic Target Database^73^, PharmGKB^74^, and the IUPHAR/BPS Guide to PHARMACOLOGY^75^. Drug–target interactions were filtered using a cutoff of binding affinities (Ki, Kd, IC50, or EC50) ≤ 10 μM to ensure high confidence in the associations following the previous research^53^. The final network comprises a total of 29,934 interactions involving 7,407 drugs. The networks in this study were visualized with Cytoscape (version 3.10.2) ^76^.

### 2.10. Network proximity-based drug repurposing

We first employed the closest network proximity method to identify potential drug repurposing candidates, as reported previously ^53,77^. The closest distance between two sets of nodes in the protein–protein interaction network, such as the drug target set (X) and a disease-related gene set (Y), was defined as follows:

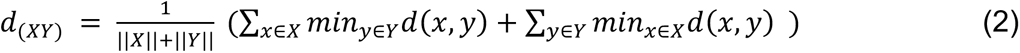

where d(x,y) represents the shortest path length between protein x in X and protein y in Y within the human protein–protein interactome. To assess the significance of the observed proximity, we calculated a Z-score for d_(X,Y)_ using a permutation-based approach. Random protein sets with similar degree distributions to X and Y were generated 1,000 times. For each random experiment, we computed the mean and standard deviation of the shortest distances. The Z-score was then calculated as:

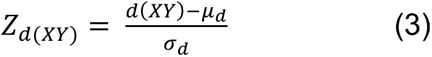

Lastly, the results were filtered using thresholds of Z-score < −2 and FDR < 0.05 to identify significant drug candidates for further investigation.

### 2.11. Gene set enrichment analysis (GSEA)

We employed a GSEA-based approach for drug repurposing, as reported previously^77,78^. Initially, drug-gene signatures were obtained from the Connectivity Map (CMap) database^79,80^, a comprehensive pharmacogenomic resource containing gene expression profiles from diverse human cell lines treated with thousands of small molecules, including both approved drugs and experimental compounds. Using the GSEA algorithm, we predicted enriched drugs for each cell type by leveraging the identified AD-associated differentially expressed genes as input. We then computed the enrichment score (ES) for each drug present in both the CMap database and the drug-target network using previously established methods^77,78^. The ES reflects the drug’s potential to reverse the gene expression patterns within the given network^81^. To ensure statistical significance and meaningful positive enrichment, the results were filtered to retain only those with a FDR < 0.05 and an ES > 0. This enabled the prioritization of candidate compounds with potential therapeutic relevance for AD.

### 2.12. Data availability

The processed data and code in this study has been submitted to the GitHub repository at https://github.com/ChengF-Lab/alzdigitaltwins.

## 3. Results

### 3.1 Study design

We obtained snRNA-seq and snATAC-seq data from 84 human brain middle temporal gyrus samples provided by the SEA-AD study^10^. Understanding and defining AD progression remains a significant challenge. Currently, AD progression is commonly assessed using neuropathologic criteria such as Braak stage, Thal phase, CERAD score, and the more comprehensive ADNC metric ^82,83^. ADNC is an integrated measure that combines multiple neuropathologic indicators, including neurofibrillary tangles (Braak), neuritic plaque scores (CERAD), and Aβ plaques(Thal)^84^. Thus, to optimally capture the distinct stages of AD progression, we categorized the 84 samples into four groups based on ADNC information from the metadata^10^: Non-AD (n = 9), Low ADNC (n = 12), Inter ADNC (n = 21), and High ADNC (n = 42) (**Fig. 1A**).

**Figure 1.**
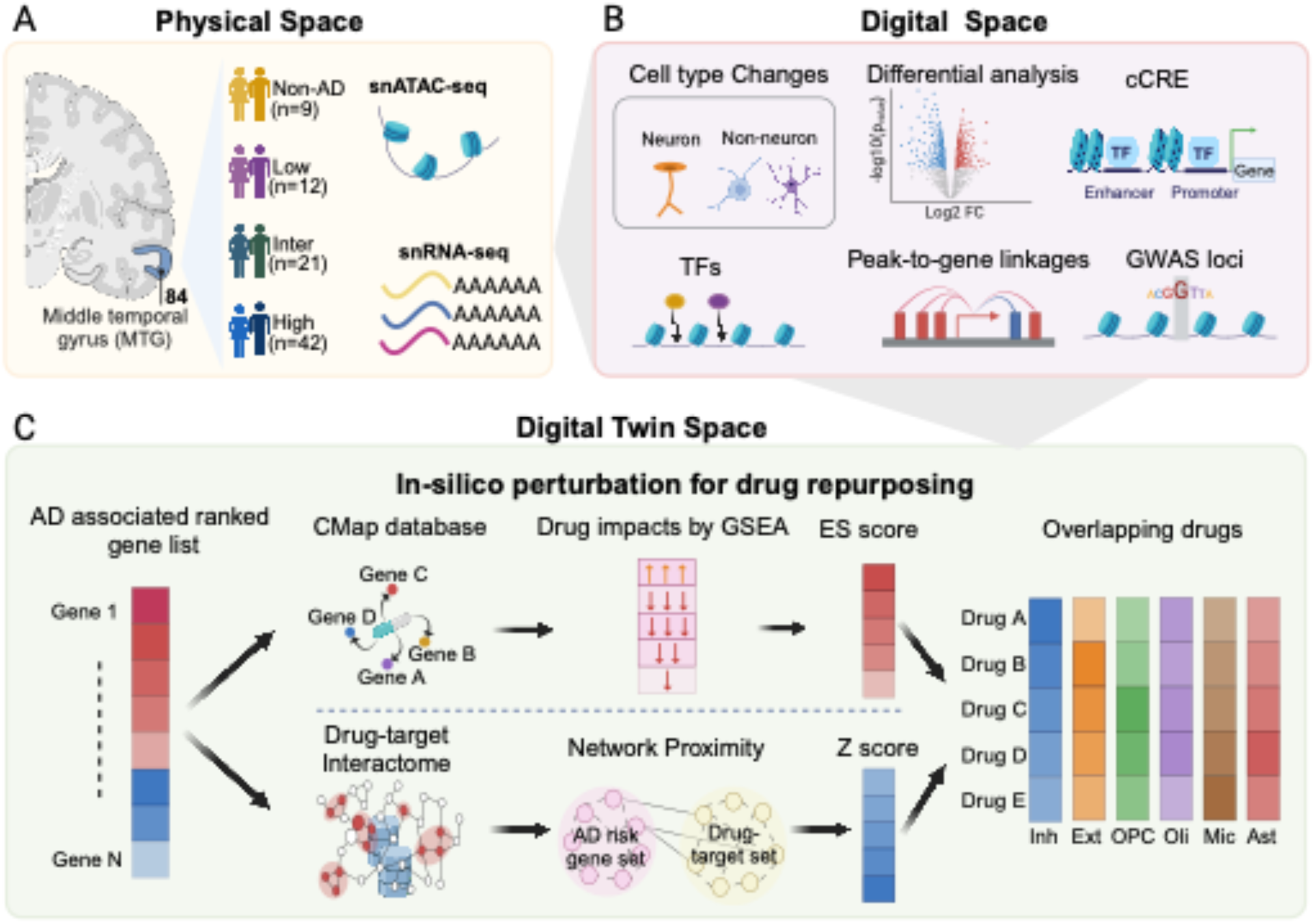
Overview of study design. (A) Collection and preprocessing of snRNA-seq and snATAC-seq datasets in physical space. (B) Multiple analysis in the digital space to identify AD associated gene list, including differential gene expression analysis, identification of cCRE, cell type-specific, AD-associated risk variants, and peak-to-gene linkages. (C) Two in-silico perturbation methods for drug repurposing within digital twin space. Network proximity analysis identifies potential drugs by assessing the proximity between drug target set and AD-related gene set. Drug efficacy is evaluated based on the proximity distance and Z-score. GSEA leverages drug-gene signatures from CMap database and differentially expressed genes to calculate enrichment score (ES). The ES reflects the drug’s potential to reverse the observed gene expression patterns within the given network.

To construct the single-cell DTs, we conducted comprehensive bioinformatics analyses that included (1) differential gene expression analysis, (2) characterization of cCREs, and (3) cell-type-specific TF-target gene networks, and (4) integrative analyses of linking peaks to genes and associating AD GWAS loci with cCREs. This enabled prioritization of ADNC-associated variants and genes within a digital framework (**Fig. 1B**). Using the AD associated gene list from digital space, we then conducted *in-silico* perturbation analyses for drug repurposing, leveraging GSEA and network proximity methods integrated with the CMap database and the drug-target interactome within a digital twin framework (**Fig. 1C**). This enabled identification of potential repurposable drugs for each cell type across AD progression.

### 3.2 Overview of snRNA-seq and snATAC-seq data across ADNC progression

AD progression staging based on ADNC does not fully align with individual neuropathologic staging metrics such as Thal phase, Braak stage, or CERAD score (**Fig. 2A**), nor with clinical cognitive status. These findings underscore the importance of defining AD progression through neuropathology to better understand the molecular changes occurring in the early stages of the disease and to inform the development of therapeutics aimed at halting AD progression. Additionally, we presented the distribution of LATE stage, apolipoprotein E 34 (*APOE4*) gene carrier status, sex, and age at death across ADNC stages in this cohort (**Fig. 2A, Supporting Information Table S1**).

**Figure 2.**
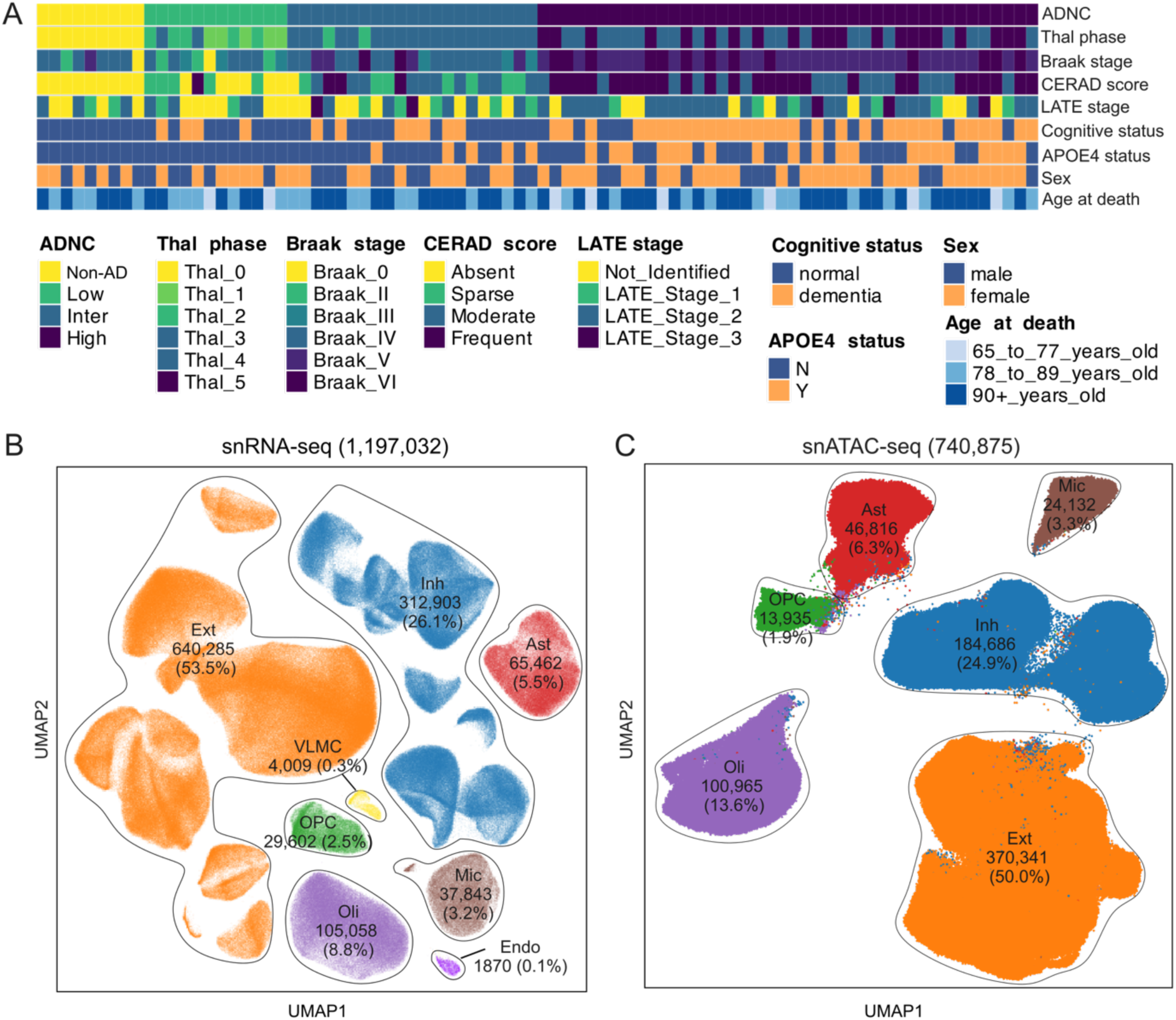
Overview of snRNA-seq and snATAC-seq datasets. (A) Heatmap displaying metadata for 84 individual samples. The rows of the heatmap, ordered from top to bottom, represent ADNC, Thal phase, Braak stage, CERAD score, LATE stage, cognitive status, APOE4 status, sex, and age at death. (B-C) UMAP representation of snRNA-seq (B) and snATAC-seq (C) are shown. The figure also shows the cell count number and proportion of each cell type.

We annotated 1,197,032 snRNA-seq cells from 84 matched individuals into 8 major cell types based on known marker genes (**Fig. 2B, Supporting Information Fig. S1A**). Inhibitory neurons (Inh; 312,903 cells, 26.1%) and excitatory neurons (Ext; 640,285 cells, 53.5%) together constitute the majority, accounting for more than half of the total cells. We further annotated 18 neuronal cell subtypes based on the classifications described in the original paper^10^ (**Supporting Information Fig. S2**). Among these, we observed that the L2/3 IT subtype of excitatory neurons showed a significant decrease of cell counts at the High ADNC stage compared to other stages, while the L4 IT subtype of excitatory neurons exhibited a significant increase at the High ADNC stage (**Supporting Information Fig. S3**). Additionally, the major glial populations included oligodendrocytes (Oli; 105,058 cells, 8.8%), oligodendrocyte progenitor cells (OPC; 29,602 cells, 2.5%), astrocytes (Ast; 65,462 cells, 5.5%), and microglia (Mic; 37,843 cells, 3.2%). Notably, microglia cell counts showed a significant increase at the High ADNC stage compared to the Non-AD and Intermediate ADNC stages (**Supporting Information Fig. S4**). Two additional cell types, endothelial cells (Endo) and vascular leptomeningeal cells (VLMC), make up a smaller fraction, with 5,879 cells (0.4%).

For snATAC-seq, we identified 740,875 cells distributed across 6 major cell types using the same marker genes as for snRNA-seq annotation (**Fig. 2C, Supporting Information Fig. S1B**). Inhibitory neurons (184,686; 24.9%) and excitatory neurons (370,341; 50.0%) showed a similar trend, comprising the largest proportions of the total cells. The remaining glial cell groups included Oli (100,965; 13.6%), OPC (13,935; 1.9%), Ast (46,816; 6.3%), and Mic (24,132; 3.3%), the cell type proportion also showed similar trends with snRNA-seq.

### 3.3 Cell type-specific gene expression change across AD progression

To identify differentially expressed genes (DEGs) across AD progression for each cell type, we conducted 6 pairwise comparisons of stage data (Low vs. Non-AD, Inter vs. Non-AD, Inter vs. Low, High vs. Non-AD, High vs. Low, High vs. Inter). A greater number of significant DEGs were observed in the High stage compared to Non-AD (5,362 DEGs), Low (4,219 DEGs), and Intermediate stages (4,216 DEGs, **Fig. 3A, Supporting Information Table S2**). We further examined gene expression changes between adjacent stages from Non-AD to High for each cell type. Notably, the DEGs from Inter to High stage showed the most pronounced gene expression changes, particularly in excitatory and inhibitory neurons, which together comprised more than half of total observed DEGs in adjacent stages (**Fig. 3B**). Additionally, among the significant DEGs identified between Inter and High stage for neuron cells, the majority were downregulated, comprising 78.64% in excitatory neurons and 82.39% in inhibitory neurons (**Fig. 3B**). Moreover, in the early stages of AD (from Non-AD to LowAD), OPC, astrocytes, and microglia exhibited significant changes, accounting for over 25% of the total DEGs. This observation indicates that there are distinct gene expression dynamics among cell types in the early stages of AD.

**Figure 3.**
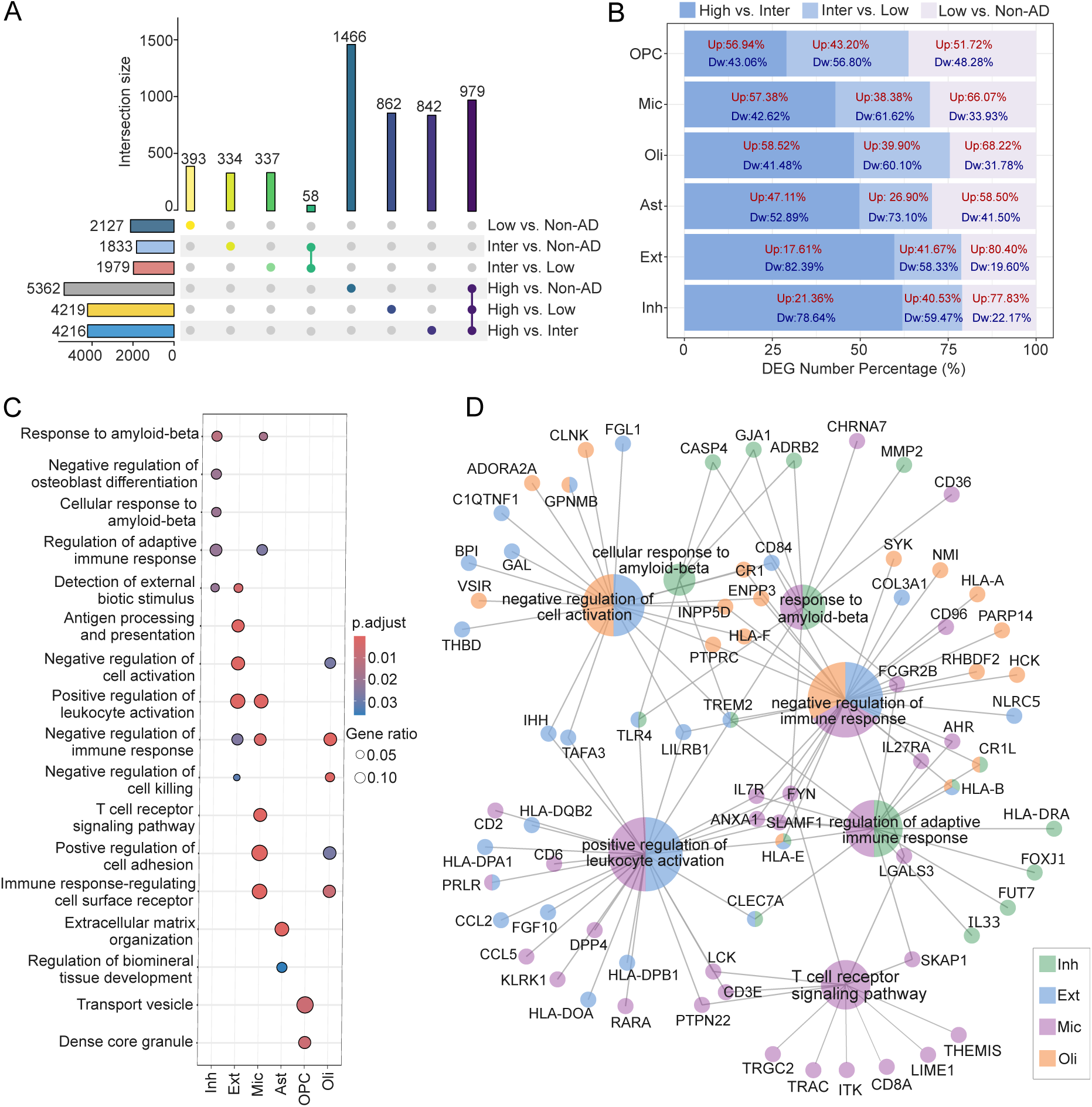
Gene expression changes across AD progression. (A) UpSet plot illustrating the number of DEGs across AD progression. The left bar plots display the total number of DEGs for each stage comparison group, while the top bar plots represent the number of DEGs that are either overlapping or unique to different groups. Colored dots connected by lines indicate the stage groups that have overlapping DEGs, while a single colored dot represents stage group without overlapping DEGs with other groups. (B) Percentage of DEGs for each cell type across adjacent stages of AD progression. Red text indicates the percentage of upregulated DEGs (Up), while blue text represents the percentage of downregulated DEGs (Dw). (C) GO function enrichment analysis of each cell for upregulated DEGs during High-vs-Inter stage. (D) Selected GO terms from (C) with corresponding genes. Different colors indicate distinct cell types, and the pie charts represent GO terms and associated genes shared across various cell types.

We further investigated the functions of genes that were up-regulated from the Inter to High stage using Gene Ontology (GO) analysis (**Fig. 3C**). We identified DEGs for each cell type that were associated with the GO terms enriched (**Fig. 3D**). Our results revealed that upregulated genes in inhibitory neurons were significantly enriched in pathways related to the response to amyloid-β, including key genes such as triggering receptor expressed on myeloid cells 2 *(TREM2)*, *ADRB2*, and *GJA1* (**Fig. 3D**). *TREM2* is involved in Aβ clearance by microglia, playing a pivotal role in mitigating Aβ plaque accumulation in AD^85,86^. Similarly, *ADRB2* is associated with signaling pathways that regulate Aβ metabolism and confer neuroprotective effects^87^, and *GJA1* contributes to mechanisms that modulate Aβ toxicity^88^. Furthermore, DEGs expressed in in excitatory neurons, microglia, oligodendrocytes and inhibitory neurons were also significantly enriched in pathways related to immune response pathways, such as *FYN, HLA-E, LCK, SYK* (**Fig. 3D**). These findings suggest that genes upregulated from the Inter to High stage are potential associated with Aβ-related processes, alongside the inflammatory and immune mechanisms underlying AD progression.

### 3.4 Identifying TF-target gene networks involved in AD progression

The accessibility of chromatin regions to TFs is a major determinant of cellular transcriptional profile^89–91^. To identify lineage-defining TFs enriched in cell type-specific accessible regions, we first defined cell type-specific peaks using the stringent criteria of Wilcoxon FDR ≤ 0.05 and Log2(fold change [FC]) ≥ 0.5, followed by TF motif enrichment analysis. For example, well-characterized TFs such as SPI1 was enriched in microglial cells as supported by footprinting evidence, whereas SOX4 and SOX9 were enriched in oligodendrocytes and JUND and FOSB were enriched in excitatory neurons (**Fig. 4A and 4B, Supporting Information Table S3**).

**Figure 4.**
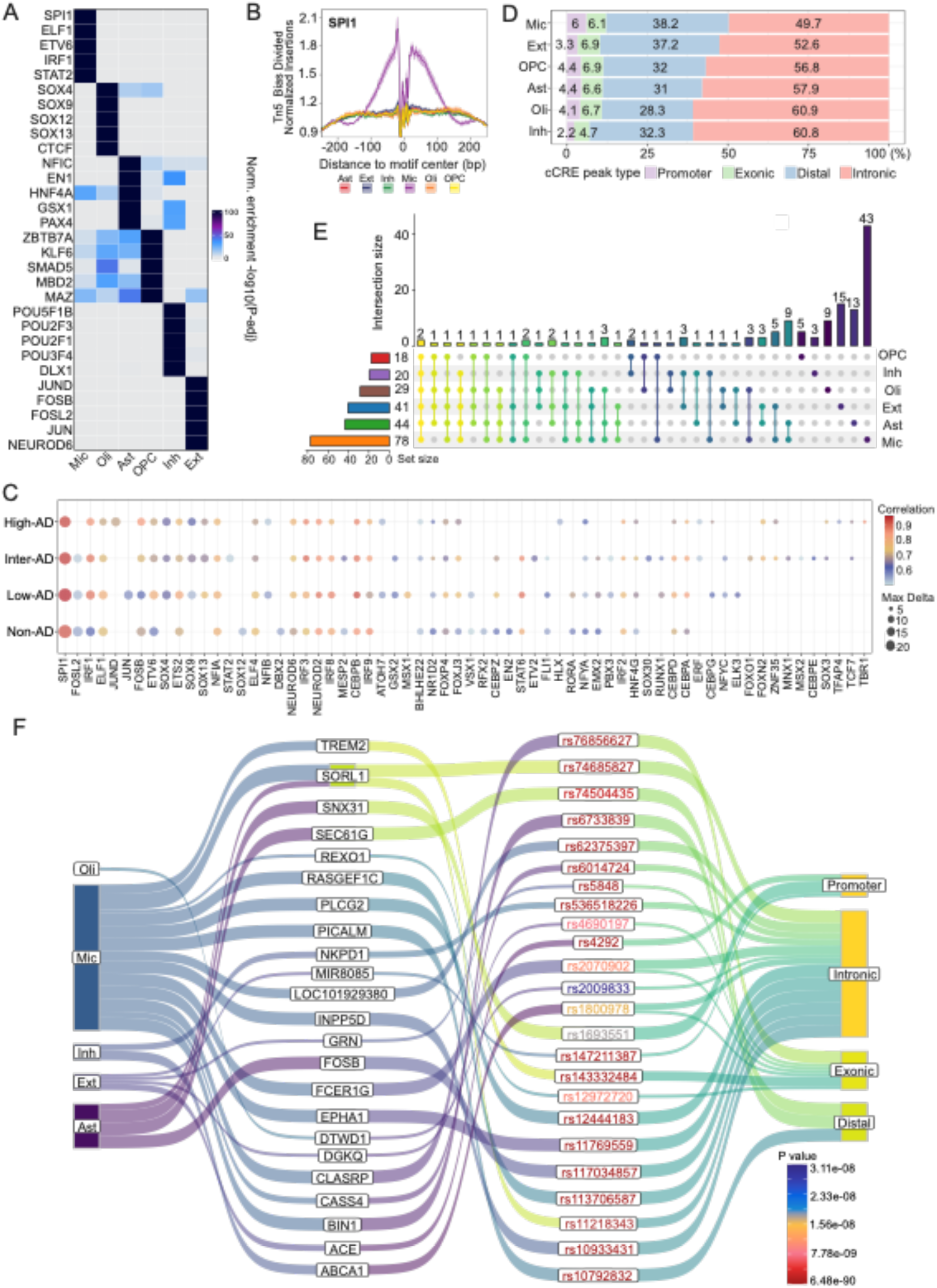
Positive TF-Regulators and cell type-specific regulatory landscapes of GWAS loci across AD progression by integrative analysis. (A) Top enriched TF motifs for each cell type. The color scale represents -log_10_ (adjusted *p*-value) of the normalized enrichment score. (B) Tn5 bias-corrected TF footprinting for SPI1. (C) Positive TF-regulators for all cell type across AD progression. (D) The percentage of genomic region annotation for cell type-specific peaks. (E) Numbers of candidate genes associated with significant AD-related SNPs for each cell type. The left bar plot shows the total number of genes per cell type, while the top bar plot highlights genes that overlap between cell types or are unique to specific cell types. (F) Sankey plot illustrating the cell type-specific AD-associated lead SNPs and their potential target genes. The first column represents cell types, the second column indicates potential target genes linked to AD-associated lead SNPs, the third column lists AD-associated lead SNPs with p-values represented by bar scale color and indicated by different text colors, and the fourth column displays genomic region annotations for peaks associated with these lead SNPs.

Some TF families share highly similar binding motifs, which can complicate identification of specific TFs associated with chromatin accessibility changes^42^. To address this challenge, we identified TFs with gene expression that correlated positively with the accessibility of their corresponding motifs by integrational analysis with snRNA-seq and snATAC-seq, designating them as “positive TF-regulators”. Across AD progression, we identified 64 positive TF-regulators based on stringent cut-off values (correlation > 0.5, adjusted *p*-value < 0.01, Z-score > 0.75) (**Fig. 4C, Supporting Information Table S4**). Specifically, the microglial-specific TF known as SPI1 was present across all stages of AD progression. In contrast, the excitatory neuron-specific TF known as JUND was predominantly enriched at the High AD stage, while STAT2 was primarily enriched at the Intermediate AD stage. These findings highlight the dynamic and cell type-specific regulatory roles of positive TF-regulators in modulating chromatin accessibility during AD progression.

### 3.5 Identifying AD progression-associated targets enriched by cCREs with AD-GWAS

We further identified cell type-specific cCREs. In total, 52,565 CREs were identified in astrocytes, 67,253 in excitatory neurons, 22,198 in inhibitory neurons, 42,241 in microglia, 25,943 in OPCs, and 24,186 in oligodendrocytes. Among these, the proportions of cCREs annotated as promoters (defined as regions 2,000 bp upstream and 100 bp downstream of a transcription start site, TSS) and distal regions (located > ±2 kb from a TSS) was much greater in microglia compared to other cell types, accounting for 6% and 38.2%, respectively (**Fig. 4D**). We also observed that excitatory neurons exhibited a comparable proportion of distal CREs accounting for 37.2%, relative to those observed in microglia (**Fig. 4D**).

We additionally identified significant GWAS loci (*p*-value < 5 × 10⁻⁸) from the GWAS summary statistics reported by Bellenguez et al.^47^. To prioritize AD-associated genes, we linked cell type-specific cCREs with AD associated SNPs by assessing overlaps between GWAS loci and peak regions. This enabled identification 141 AD-associated genes (**Supporting Information Table S5**). Of these, 78 genes were associated with microglia, including 43 genes that were specific to microglia, including such as *FCER1G, APOC4-APOC2, INPP5D* (**Fig. 4E**). Additionally, 44 AD-associated genes were linked to astrocytes, with 13 genes being astrocytes-specific, such as *SEC61G, CLU*, and 41 genes were associated with excitatory neurons including 15 genes that were excitatory neuron-specific, such as *DGKQ, GRN* (**Fig. 4E**).

Next, we retrieved 88 lead SNPs directly from Bellenguez et al^47^ and linked these lead SNPs to cCREs (See Methods, **Supporting Information Table S6**). This analysis further highlighted 23 AD-associated genes enriched in a cell type-specific manner (**Fig. 4F**). Specifically, we identified 13 AD-associated genes in microglia, including well-known AD genes, such as *TREM2* and *BIN1*. *TREM2* was associated with the rs143332484 locus (AD GWAS *p*-value: 6.035 x 10^-19^), with its corresponding CRE mapped to an exonic region, whereas *BIN1* was linked to the rs6733839 locus (AD GWAS *p*-value: 6.48 x 10^-90^), with its CRE located in a distal regulatory region (**Fig. 4F**). Additionally, AD-associated genes such as *ACE*, *DGKQ*, *GRN*, and *MIR8085* were enriched in excitatory neurons, and genes such as *SNX31*, *SEC61G*, and *FOSB* were observed in astrocytes. Furthermore, *ABCA1* and *NKPD1* were associated with inhibitory neurons, and *DTWD1* was linked to oligodendrocytes. Collectively, we identified and prioritized cell type-specific AD-associated genes through integrating regulatory landscapes associated with cCREs and AD-GWAS data across diverse cellular populations.

### 3.6 Cell type-specific TF regulatory network in AD progression

To gain further insight into TF-mediated gene regulation across AD progression, we constructed cell type-specific positive TF regulatory networks. We identified candidate target genes for positive TF-regulators by assessing the correlation between motif activity and gene expression, as well as the linkage score for each gene-TF pair (**Supporting Information Table S6**). We selected several TFs specific to microglia (SPI1, IRF1, and CEBPB) (**Fig. 5A**), excitatory neurons (JUN, JUND, FOSB, NEUROD2, and NEUROD6) (**Fig. 5B**), and oligodendrocytes (SOX4, SOX9, SOX12, and SOX30, **Fig. 5C**) to construct the TF-target networks. Within these networks, we highlighted TF targets that are annotated as DEGs across AD progression and genes located at AD GWAS loci. For instance, *APOE* as the candidate target of CEBPB is a well-established AD-associated gene that was also found to have significantly upregulated DEGs during the transition from the Inter to High stage of AD. Additionally, it is located within an AD-associated GWAS locus (**Fig. 5A**). These findings underscore the pivotal role of cell type-specific TFs in regulating key genes associated with AD progression, linking transcriptional changes to genetic risk loci and highlighting potential mechanisms underlying disease pathology.

**Figure 5.**
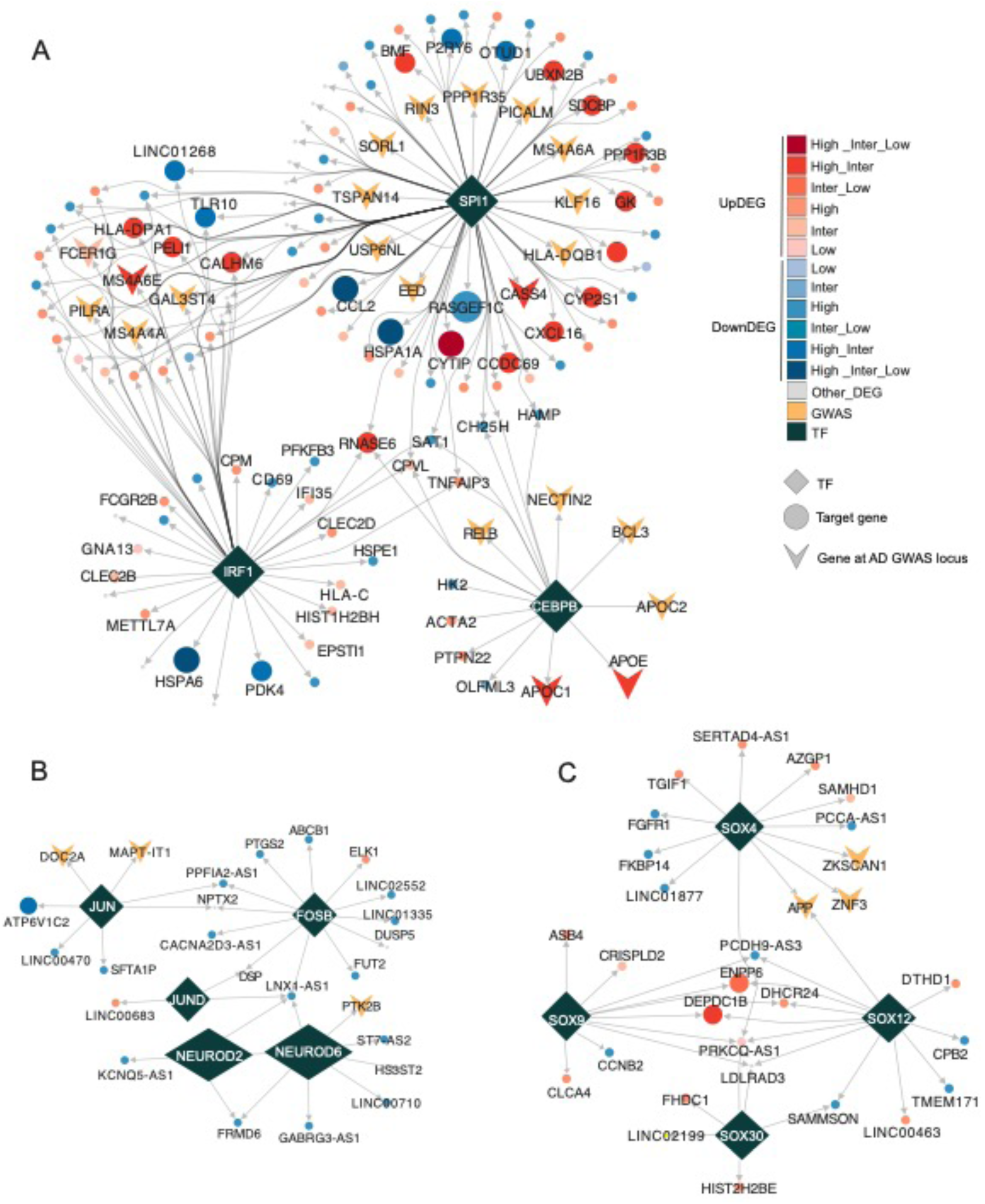
Cell type-specific TF regulatory network. (A) TF regulatory networks in microglia for SPI1, IRF1, and CEBPB, highlighting predicted candidate target genes that exhibit significant expression change during AD progression or are annotated in AD-asscoiated GWAS locus. (B) TF regulatory networks in excitatory neurons for the TFs JUN, JUND, FOSB, NEUROD2, and NEUROD6, showing the predicted candidate target genes that significantly changed during AD progression or annotated in AD GWAS locus. (C) TF regulatory networks in oligodendrocytes for the TFs SOX4, SOX9, SOX12, and SOX30, showing the predicted candidate target genes that significantly changed during AD progression or annotated in AD GWAS locus. In all panels, the shape of diamonds represents TFs, and circle represents target genes of TFs, and “V” arrows indicate the target genes of TFs located within AD associated GWAS loci. The red gradient denotes target genes annotated with upregulated DEGs at different stages of AD progression, while the blue gradient represents target genes annotated with downregulated DEGs. Genes with inconsistent DEG change across stages are labeled in gray.

### 3.7 Discovery of cell type-specific repurposable drugs for AD progression

In total, we identified 141 potential AD-associated genes based on prior analyses (**Fig. 6A**). Among these, 23 genes were associated with AD lead SNPs (e.g., *ACE, TREM2, B1N1*), 48 genes were associated with significant DEGs across AD progression (e.g., *ABCA1, APOE, APOC1, APOC2*), 52 genes were target genes of positive TF-regulators (TG) (e.g., *DGKQ, APOE, SNX31*), and 36 genes (e.g., *ACE, APP, APOE, ABCA1, CRHR1*) were known drug targets with FDA-approved or clinically investigational therapies (**Fig. 6A**).

**Figure 6.**
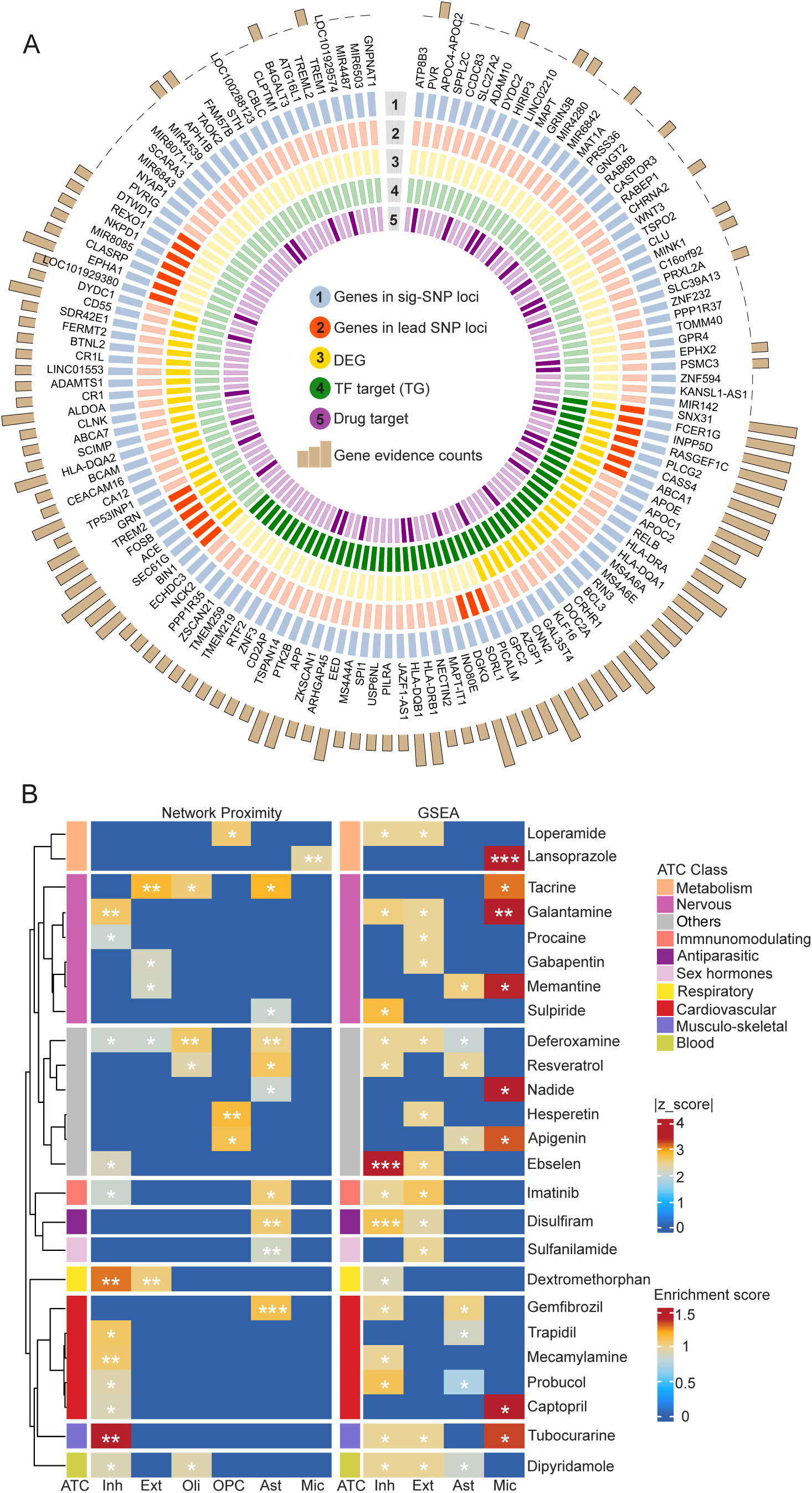
Repurposable drug candidates for cell type-specific across AD progression. (A) Prioritized AD-associated genes by combining cCREs and significant AD associated variants. The first circle with blue rectangles highlights genes identified by overlapping AD-associated significant SNPs loci with cell type-specific cCREs. The second circle with dark red rectangles represents genes associated with AD lead SNPs loci and cCREs. The third circle with dark yellow rectangles is DEGs. The fourth circle with dark green rectangles represents the target genes regulated by TFs. The fifth circle with dark purple rectangles comprises target genes annotated as known drug targets. And the outer bar plot provides the evidence counts for each gene. (B) Predicted repurposable drug candidates by network proximity and GSEA clustered by ATC first level class. Left panel is enriched drug candidates identified by network proximity method, where gradient colors reflect absolute Z-scores. Right panel is enriched drug candidates identified by GSEA method, where gradient colors represent enrichment scores. Statistical significance is denoted by * (FDR < 0.05), ** (FDR<0.01), *** (FDR < 0.001).

To predict cell type-specific drug candidates across AD progression, we utilized 141 potential AD-associated genes as input for a network proximity analysis. This approach evaluates the network relationship between AD-associated genes and drug targets within the human PPI network. We subsequently identified enriched drug candidates by performing GSEA integrating drug-gene signatures from the CMap database and information from the DEGs (see Methods). The ES indicates each drug’s potential to reverse the observed gene expression patterns. By integrating these methods, we identified 25 overlapping candidate drugs that met the criteria for both network proximity (Z-scores < −2, FDR < 0.05) and GSEA (ES > 0, FDR < 0.05) (**Fig. 6B, Supporting Information Table S7**). These 25 drugs were categorized into 10 pharmacological classes based on the first level of the Anatomical Therapeutic Chemical (ATC) classification system, with the most candidate drugs targeting the nervous and cardiovascular systems.

Among these, 13 drugs were consistently identified across cell types by both methods, including galantamine, mecamylamine, dextromethorphan, tubocurarine, gabapentin, ebselen, resveratrol, imatinib, dipyridamole, deferoxamine, gemfibrozil, lansoprazole, and probucol. Galantamine, which emerged as a potential drug candidate in inhibitory neurons, excitatory neurons, and microglia, is an acetylcholinesterase inhibitor approved for the treating mild to moderate AD^92^. By inhibiting acetylcholine breakdown and modulating nicotinic receptors, it enhances cholinergic transmission, thereby improving cognitive function. However, Galantamine does not halt AD progression^93^.

Beyond galantamine, other identified drugs have also shown promise in preclinical studies. For instance, ebselen, identified as a potential drug candidate in inhibitory and excitatory neurons, is a synthetic organoselenium compound with antioxidant and anti-inflammatory properties^94^. By mimicking glutathione peroxidase activity, ebselen reduces oxidative stress, which is implicated in AD pathology. Similarly, Resveratrol, highlighted as a potential drug candidate in astrocytes, is a polyphenolic compound found in grapes and red wine with antioxidant properties^95–97^. Notably, it has been extensively studied for its ability to modulate amyloid-beta aggregation and reduce neuroinflammation in AD^95–98^, both critical in AD pathology.

Another promising candidate, Imatinib, was identified in inhibitory and excitatory neurons. Known for its ability to modulate gamma-secretase activity^99,100^, Imatinib has been investigated for its potential to reduce amyloid-beta production. Additionally, gabapentin, identified in excitatory neurons, is an anticonvulsant that modulates voltage-gated calcium channels that has been studied for off-label use in managing behavioral symptoms associated with dementia, such as agitation^101^.

To provide deeper insights into the mechanism-of-action of the drug candidates, we constructed drug-target networks for the 13 prioritized drugs (**Fig. 7**) and an additional 12 candidate drugs (**Supporting Information, Fig. S5**). For example, galantamine was found to target multiple subunits of the nicotinic cholinergic receptor — *CHRNA2, CHRNA10, CHRNB1, and CHRND*, thereby potentially enhancing cholinergic signaling, a critically impaired pathway in AD^102^. Similarly, mecamylamine and tubocurarine interact with specific nicotinic cholinergic receptor subunits, including *CHRNA7* and *CHRNA2* (**Fig. 7**), further emphasizing the role of these receptors in AD pathophysiology. Furthermore, resveratrol and deferoxamine target *APP*, a key gene involved in the pathophysiology of AD^103^. These findings highlight the potential of drug candidates to modulate critical pathways in AD progression in a cell type-specific manner. However, more functional and clinical observations are highly warranted for those top predicted candidate drugs in the future.

**Figure 7.**
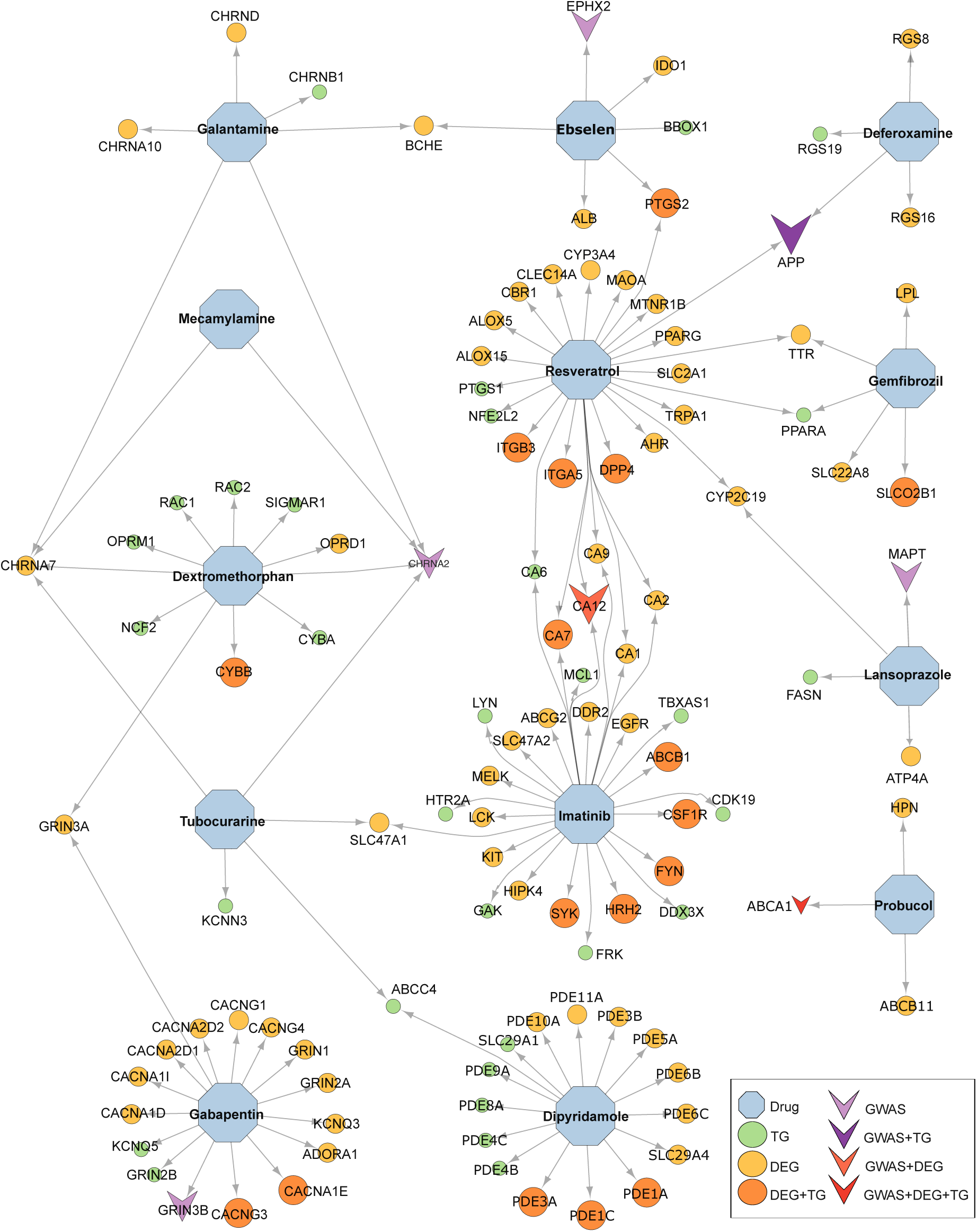
Network of targets for 13 potential drug candidates. This network illustrates the relationships between the 13 potential drug candidates and their respective targets. The shape of hexagon represents the drug, and circle represents drug targets. “V” arrow denotes additional annotations for drug target genes that are located within the GWAS loci. Green circles indicate drug target genes annotated as TGs of TFs. Yellow circles represent drug target genes identified as DEGs, and orange circles denote the drug target genes annotated as both DEGs and TGs of TFs. Light purple arrows represent drug target genes located within AD-associated GWAS loci, dark purple arrows indicate drug target genes located within AD-associated GWAS loci and annotated as TGs. Orange arrows highlight drug target genes located within AD-associated GWAS loci and annotated as DEGs, and red arrows represent drug target genes located within AD-associated GWAS loci and annotated as both TGs and DEGs.

## 4. Discussion

Despite significant efforts to discover effective drugs for AD, the complex neuropathological mechanisms, cellular heterogeneity, and challenges associated with early diagnosis continue to hinder drug development^1,2,98,104^. To address these challenges, we have introduced here the concept of single-cell digital twin with a comprehensive scDT framework that integrates a network-based approach with snRNA-seq and snATAC-seq data for therapeutic target identification and drug repurposing in AD progression. Our framework categorizes AD progression into four distinct stages based on ADNC that is a robust classification of neuropathology. This dynamic staging allows for detailed observation of cellular and molecular changes during disease progression, enabling identification of precise therapeutic targets to inform drug development.

Through this approach, we observed cell type-specific transcriptomic dynamics, with particularly pronounced changes for each cell in gene expression from the intermediate to high stages of AD. These findings highlight critical molecular shifts in gene regulation that are associated with late stages of AD, consistent with previous studies^5^. Notably, in the early stages of AD (from Non-AD to Low-AD), OPC, astrocytes, and microglia exhibited significant changes, contributing to more than 25% of the total DEGs. This indicates that gene expression dynamics vary distinctly among cell types during AD progression. These results collectively emphasize the heterogeneous nature of transcriptional changes across cell types during AD progression.

We also identified cell-specific CREs and positive TFs regulatory networks, which were analyzed in conjunction with AD associated GWAS. This integrative approach revealed 141 AD associated genes, including 36 known targets of FDA-approved or clinically investigational therapies. Among these, several are implicated in amyloid pathology and tau dynamics, such as *APP, ACE* and *MAPT*, which are central to amyloid-beta processing and tau pathology, both hallmark features of AD^105^. *ADAM10* contributes to the non-amyloidogenic cleavage of *APP*, reducing amyloid-beta production^106^. Several genes involved in lipid metabolism and cholesterol transport, including *APOE, ABCA1, and CLU*, are critical in lipid handling and amyloid-beta clearance^107–109^, with additional contributions from *AZGP1* and *DGKQ*^47,110^. Furthermore, neuroinflammation and immune modulation were linked to many genes including *TREM1, INPP5D, HLA-DRA, HLA-DRB1, HLA-DQB1, HLA-DQA2, FCER1G*, and *CRHR1*^111–114^. Novel and emerging targets, such as *LINC02210, PRXL2A, EED, B4GALT3, MINK1, CD55, and AOK2*, were also identified. This comprehensive analysis underscores the diverse molecular mechanisms contributing to AD pathology and highlights promising therapeutic targets for future drug development.

Leveraging these 141 AD-associated genes, we further identified 25 candidate drugs by network proximity and GSEA analyses, 13 of which were consistently identified by both methods. These drugs exhibit diverse mechanisms of action. For instance, mecamylamine, a nicotinic acetylcholine receptor antagonist, regulates neuronal excitability and reduce neuroinflammation^115,116^. Ebselen targets proteins such as EPHX2, IDO1, and PTGS2 (**Fig. 7**). EPHX2 encodes soluble epoxide hydrolase (sEH), an enzyme involved in converting epoxides into diols. These lipid signaling molecules play crucial roles in regulating inflammation, vascular function, and oxidative stress^117–120^. *IDO1* and *PTGS2* are key regulators of immune responses and inflammation^121,122^, suggesting that ebselen may impact pathways linked to inflammation, oxidative stress, and metabolic regulation. Dipyridamole mainly targets phosphodiesterase (PDE) genes, such as *PDE1A, PDE3A, PDE1C, PDE5A*, which regulate intracellular levels of cyclic nucleotides like cAMP and cGMP—critical mediators of neuronal signaling, plasticity, and memory^123^. Imatinib interacts with multiple proteins, including ABCB1, ABCG2, SYK, LCK, FYN, KIT, DDR2, CA7, and CA12 (**Fig. 7**). Notably, *ABCB1* and *ABCG2* are associated with amyloid-beta clearance and brain detoxification^124,125^. SYK serves as a critical regulator of microglial activation^126^, while FYN and KIT play essential roles in synaptic function^127,128^. Gemfibrozil is a lipid-lowering drug that acts on PPARA, LPL, SLCO2B1, SLC22A8, and TTR^129^. While lansoprazole is primarily used for gastric acid suppression, its potential effects on target genes like *MAPT, FASN, and ATP4A,* suggesting possible connections to pathways relevant to AD^130–132^. Probucol’s modulation of ABCA1, ABCB11, and HPN highlights its potential for addressing lipid dysregulation, amyloid-beta clearance, and neuroinflammation in AD^133,134^.

Despite the promising findings, we acknowledge several limitations in this study. A significant constraint is the relatively small number of brains from which the original tissue was derived, which may limit the generalizability of our results. Additionally, further functional and clinical validation of the top predicted drug candidates is crucial to gain deeper insights into their mechanisms of action and therapeutic efficacy. It is also important to note that some candidate therapies, such as resveratrol, have previously failed in clinical trials, emphasizing the need for additional research to bridge the molecular insights presented here with their potential success in therapeutic applications. To address these limitations, we plan to incorporate more single-cell multi-omics data from the Religious Orders Study and Memory and Aging Project (ROSMAP) and leverage generative AI methods to develop a comprehensive single-cell foundation model. This model aims to accurately represent AD progression and further refine the single-cell digital twin framework, paving the way for more robust therapeutic target identification and drug development.

## 5. Conclusions

In summary, we developed a single-cell digital twin framework that integrates snRNA-seq and snATAC-seq data to elucidate cell type-specific CREs and positive TF-regulators networks associated with AD progression. This framework enabled us to prioritize 141 AD-associated genes and identify 13 candidate drugs for key cell types using a combination of network proximity analysis and GSEA methods. This provides new insights into the molecular mechanisms and therapeutic opportunities across the trajectory of AD progression.

## Data and code availability

Raw snRNA-seq and snATAC-seq data used from this study have been deposited in the AD Knowledge Portal (syn26223298, https://adknowledgeportal.synapse.org/Explore/Studies/DetailsPage/StudyDetails?Stud y=syn26223298). Code used for all data analysis are available in the GitHub repository: https://github.com/ChengF-Lab/AD-digitaltwins.

## Author contributions

F.C. conceived the study. Y.R. designed and performed all data analyses and experiments. M.H., Y.E.L., A.A.P., and J.C. interpreted the data analysis. J.C. provided clinical context of the outcomes. Y.R. drafted the manuscript. Y.R., F.C., A.A.P. and J.C. critically revised the manuscript. All authors gave final approval of the manuscript.

## Consent Statement

The consent was not necessary for this study.

